# Control of Cucumber green mottle mosaic virus in commercial greenhouse production with agricultural disinfectants and resistant cucumber varieties

**DOI:** 10.1101/2020.06.24.168302

**Authors:** W. Ellouze, V. Mishra, R. J. Howard, K.-S. Ling, W. Zhang

**Author notes:** Corresponding author: Dr. Walid Ellouze (Address: Agriculture and Agri-Food Canada, 4902 Victoria Avenue North, Vineland Station, ON, L0R 2E0, Canada;). Emails of authors: Dr. Walid Ellouze, Dr. Vachaspati Mishra, Dr. Ronald J. Howard, Dr. Kai-Shu Ling, Dr. Weizheng Zhang.

## Abstract

Cucumber green mottle mosaic virus (CGMMV) is a re-emerging threat to greenhouse cucumber and other *Cucurbitaceae* crop productions worldwide. This seed-borne virus can easily spread from a contaminated seed to seedlings and to adjacent plants through mechanical contact of the foliage of diseased and healthy plants causing extensive yield losses. Additionally, infection may not be limited to the current crop but may also affect subsequent crops due to the long-term persistence of the virus on contaminated crop residues, greenhouse hard surfaces and soil or soil-less greenhouse substrates. In the present work, three greenhouse trials were conducted to develop an integrated pest management strategy towards controlling CGMMV in commercial cucumber greenhouses, by implementing an effective sanitization program and using resistant and grafted cucumber varieties. Results of sanitization trial highlighted that pressure washing and cleansing with an alkaline foam cleanser has eliminated CGMMV on some of the most heavily infested areas. However, three successive applications of cleanser and disinfectants were essential to completely eliminate CGMMV on porous and uneven surfaces, such as cement alleyway, tray gutter and floor mats. The varietal trial revealed that out of 15 cucumber varieties evaluated, two Mini (‘Katrina’ and ‘Khassib’) and three Long English (‘Sepire’, ‘Bomber’ and ‘LC13900’) had reduced or delayed CGMMV infection spread in the greenhouse but were intermediate in yield. The varieties ‘Sunniwell’ and ‘Bonbon’ were the most tolerant to CGMMV. They showed a high CGMMV infection level without compromising yield. These results proved the need for new productive cucumber varieties with CGMMV resistance. Grafting experiment showed only yield increase in case of grafted ‘Picowell’ over ‘Bonbon’ and not marked CGMMV resistance, which is a much desirable result when the grafting experiments are evaluated for their economic potential. In all, the current experimental trials unfold unique methodologies on CGMMV management in commercial greenhouses that are recommended to the growers to be followed for reducing crop losses and get benefitted on revenue compromise.

## Introduction

Cucumber green mottle mosaic virus (CGMMV), a member of the genus *Tobamovirus* in the family *Virgaviridae*, is an increasing threat to cucumber (*Cucumis sativus* L.) and other *Cucurbitaceae* crop productions globally (Dombrovsky *et al.*, 2017; Mandal *et al.*, 2008). This re-emerging virus can cause extensive damage to cucumber crops resulting in substantial yield losses and a lower market value (Dombrovsky *et al.*, 2017; Fletcher *et al.*, 1969). This seed-borne virus can easily spread across short and long distances through the use of contaminated seeds and infected seedlings (Li *et al.*, 2016; Liu *et al.*, 2014; Sui *et al.*, 2019). CGMMV was first described in 1935 in the United Kingdom (Ainsworth, 1935), and eventually it spread worldwide to almost all cucurbit-producing regions (Al-Shahwan and Abdalla, 1992; Borodynko-Filas *et al.*, 2017; Budzanivska *et al.*, 2007; Celix *et al.*, 1996; Dombrovsky *et al.*, 2017; Kim *et al.*, 2010; Ling *et al.*, 2014; Moradi and Jafarpour, 2011; Reingold *et al.*, 2013; Tesoriero *et al.*, 2016; Tian *et al.*, 2014; Varveri *et al.*, 2002; Zhang *et al.*, 2009). CGMMV can infect a number of common weed species that includes *Euphorbiaceae, Solanaceae, Lamiaceae, Boraginaceae, Apiaceae, Amaranthaceae, Chenopodiaceae* and *Portulacaceae*, which in turn may serve as a virus reservoirs (Boubourakas *et al.*, 2004; Dombrovsky *et al.*, 2017; Shargil *et al.*, 2017).

Although the level of natural virus transmission through seed is relatively low (Lecoq and Desbiez, 2012), the ease of mechanical transmission of CGMMV from a contaminated seed to seedlings and to adjacent plants, especially in propagation houses, makes this virus very contagious (Sui *et al.*, 2019). The mechanical spread of the virus may occur by various means, including the handling of plants, leaf contact, wounds made with cutting tools, by farm equipment (Li *et al.*, 2016), chewing insects such as the cucumber leaf beetle (*Raphidopalpa fevicollis*) (Rao and Varma, 1984), and pollinators such as European honeybees (*Apis mellifera* L.) (Darzi *et al.*, 2018). The presence of a single CGMMV-infected plant in a cucumber greenhouse may result in the eventual infection of the entire crop. In addition, the virus is extremely stable and its particles may remain viable for several months in crop residues, soil and on greenhouse hard surfaces under relatively extreme climatic conditions (Varveri *et al.*, 2002). This property of stability, combined with its high infectivity rate through mechanical contact with the foliage, and capacity to affect subsequent greenhouse crops, have increased the economic importance of this virus. High CGMMV infections may force growers to terminate their crops early because of unproductiveness, hence reducing the overall profitability of their operations (Fletcher *et al.*, 1969; Reingold *et al.*, 2016).

To date, no effective chemical or cultural control methods have been developed to prevent CGMMV spread. Multiple approaches should be developed and adopted to prevent the introduction and delay the spread of the virus in commercial cucumber greenhouses. Using an integrated pest management approach will minimize the negative effects of CGMMV on cucumber production and keep the disease damage under the economic threshold. CGMMV disease management must include removal of virus reservoirs, phytosanitary practices, use of certified virus-free seed, selection of a resistant cultivars, and the use of grafted plants (Ellouze *et al.*, 2015).

Greenhouse sanitization, which consists of cleaning and disinfection, is the cornerstone of an effective greenhouse integrated pest management (IPM)/Biosecurity program. It is an effective process of decontaminating surfaces that may have become contaminated with pathogens, insects, mites, nematodes, weeds, algae, etc. An effective sanitization program should lead to the reduction or elimination of active and dormant stages of pathogens and pests, as well as to disrupt their life cycle. Cleaning a commercial greenhouse facility after infection of a cucumber crop with CGMMV is a difficult task and requires patience and attention to detail because the virus is persistent on the greenhouse structure itself, as well as on substrate bags, walkways, bench tops, troughs, small equipment, produce baskets, workers’ clothing and hands, and on many other surfaces within production facilities. Therefore, understanding where the virus can be found in a commercial greenhouse and how persistent it is in the environment is a key step in developing an effective sanitization program. Effective sanitization strategies for this virus are mainly aimed at reducing or eliminating existing sources of infection and the prevention of virus transmission.

Host resistance is one of the most desirable viral disease management strategies and some commercially available greenhouse cucumber cultivars are described by the seed companies as having high or intermediate resistance to CGMMV (Table 1). However, there is limited knowledge regarding the genetic mechanisms involved in resistance to CGMMV in cucumbers and other cucurbits. Two partially resistant *C. sativus* accessions were identified with mild CGMMV disease symptoms (Crespo *et al.*, 2018). Although, no commercially available greenhouse cucumber varieties are immune to CGMMV, a better understanding of their relative resistance and susceptibility to the Canadian isolate of CGMMV would be beneficial. Science-based recommendations need to be available to greenhouse cucumber growers, including the opportunity to select the most disease resistant/tolerant and agronomically suitable varieties for CGMMV management in commercial greenhouses.

**Table 1.** List of Mini and Long English cucumber varieties evaluated in trial and their level of resistance to different diseases according to the seed companies.

| Cucumber Type | Seed Company | Variety Name and (#) | High Resistance | Intermediate Resistance |
| --- | --- | --- | --- | --- |
| Mini | Rijk Zwaan | Sunniwell (1) | Ccu | **CGMMV/CMV/CVYV/Px** |
|  |  | Deltastar (2) | Cca/Ccu/Px | CMV/CVYV |
|  |  | RZ 22-551 (3) | **CGMMV** | - |
|  |  | Khassib (4) | Ccu/Px | CMV/CVYV/PRSV/WMV/ZYMV |
|  | Monsanto/De Ruiter | Jawell (5) | - | CMV/CVYV/Px |
|  | Enza Zaden | Katrina (6) | Ccu | Px/CMV/CVYV |
| Long English | Monsanto / De Ruiter | DR4879CE (7) | Px | **CGMMV** |
|  | Syngenta | Bomber (8) | Px | **CGMMV/CMV** |
|  |  | LC13900 (9) | **CGMMV** | - |
|  | Enza Zaden | Dee Lite (10) | Ccu | **CGMMV/CMV/CVYV/Px** |
|  |  | Komet (11) | Cca/Ccu | **CGMMV/CMV/CVYV/Px** |
|  | Rijk Zwaan | Bonbon (12) | **CGMMV/Ccu** | CVYV/Px |
|  |  | Verdon (13) | **CGMMV/Cca/Ccu/Px** | CMV/CVYV |
|  |  | Addison (14) | Ccu/Px | **CGMMV/CMV/CVYV** |
|  | Nunhems | Sepire (15) | **CGMMV** | - |

Cca - *Corynespora* leaf spot caused by *Corynespora cassiicola*; Ccu - Scab and gummosis caused by *Cladosporium cucumerinum*; CVYV - Cucumber vein yellowing virus; CGMMV - Cucumber green mottle mosaic virus; Px - Powdery mildew caused by *Podosphaera xanthii*; WMV - Watermelon mosaic virus; PRSV - Papaya ringspot virus; ZYMV - Zucchini yellow mosaic virus; CMV - Cucumber mosaic virus.

While breeding for viral disease resistance is a long-term process, vegetable grafting offers a short term alternative for compiling tolerance traits and improving yield potential (Pérez-Alfocea, 2015; Rouphael *et al.*, 2018). Grafted plants have greater tolerance to biotic and abiotic stresses thus providing higher fruit yields (Louws *et al.*, 2010; Melnyk, 2016). The enhanced tolerance in grafted plants has been attributed to increases in vigor, improved photosynthetic efficiency, stronger antioxidative defense system, heightened hormonal signaling and long-distance movement of mRNAs, small RNAs and proteins (Davis *et al.*, 2008; Rouphael *et al.*, 2018). Grafting on resistant rootstocks has been reported to confer resistance in cucumber to foliar fungal pathogens, such as target leaf spot, powdery mildew and downy mildew (Gu *et al.*, 2008; Guan *et al.*, 2012; Hasama *et al.*, 1993; Sakata *et al.*, 2006; Shibuya *et al.*, 2015). Tolerance to viruses in seedless watermelon plants was reported to be improved by grafting (Wang *et al.*, 2002). *Pepino mosaic virus, Tomato yellow leaf curl virus* and *Tomato spotted wilt virus* were also reported to be controlled by grafting (Louws *et al.*, 2010). In hydroponic production systems, roots of a disease-resistant rootstocks provide an extra line of defense against pathogens in potentially contaminated recirculating nutrient solution. Currently, knowledge about CGMMV resistant rootstocks is limited, and information on the usage of plants grafted onto resistant rootstocks in commercial greenhouses is not readily available.

The objective of our study was to develop a management strategy towards controlling CGMMV in commercial cucumber greenhouses by implementing an effective sanitization program and using virus-resistant and grafted cucumber varieties to effectively manage CGMMV disease without compromising overall crop production and impacting its economic value.

## Materials and methods

### Evaluation of the effectiveness of the different sanitization steps

An extensive environmental sampling program of hard surfaces within a 15-acre commercial cucumber greenhouse with a high incidence of CGMMV disease (>50% infected plants) in Alberta, Canada was undertaken before and after each of the four sanitization steps being used by the grower (Fig. 1). Before crop removal, three of the most diseased greenhouse areas (>90% infected plants) were identified. Solar-Cult^®^ Pre-moistened Sampling Cellulose Sponge Kit (www.solarbiologicals.com) was used for the collection of samples from 15 different hard surfaces in each of the three key infested areas of the operation that included: cement alleyways, tray gutters, rail brackets, rails, floor fabrics, water hoses below trays, tray end irrigation/drain hoses, canopy heating pipes, tray tops, support posts, steel beam tray supports, cropping wires, perimeter heating pipes, interior walls, and shade curtains on walls. The samples were collected post-crop removal, post pressure washing and after cleansing with an alkaline foam cleaner (MS Topfoam LC ALK, Schippers Canada Ltd., Alberta, Canada), and post application of alkaline (C-Clean, Ontario, Canada) and peroxide surface disinfectants (Virkon^®^Greenhouse, Vétoquinol N.-A. Inc., Quebec, Canada) separately, according to the manufacturer’s instructions (Fig. 1).

**Fig. 1.**
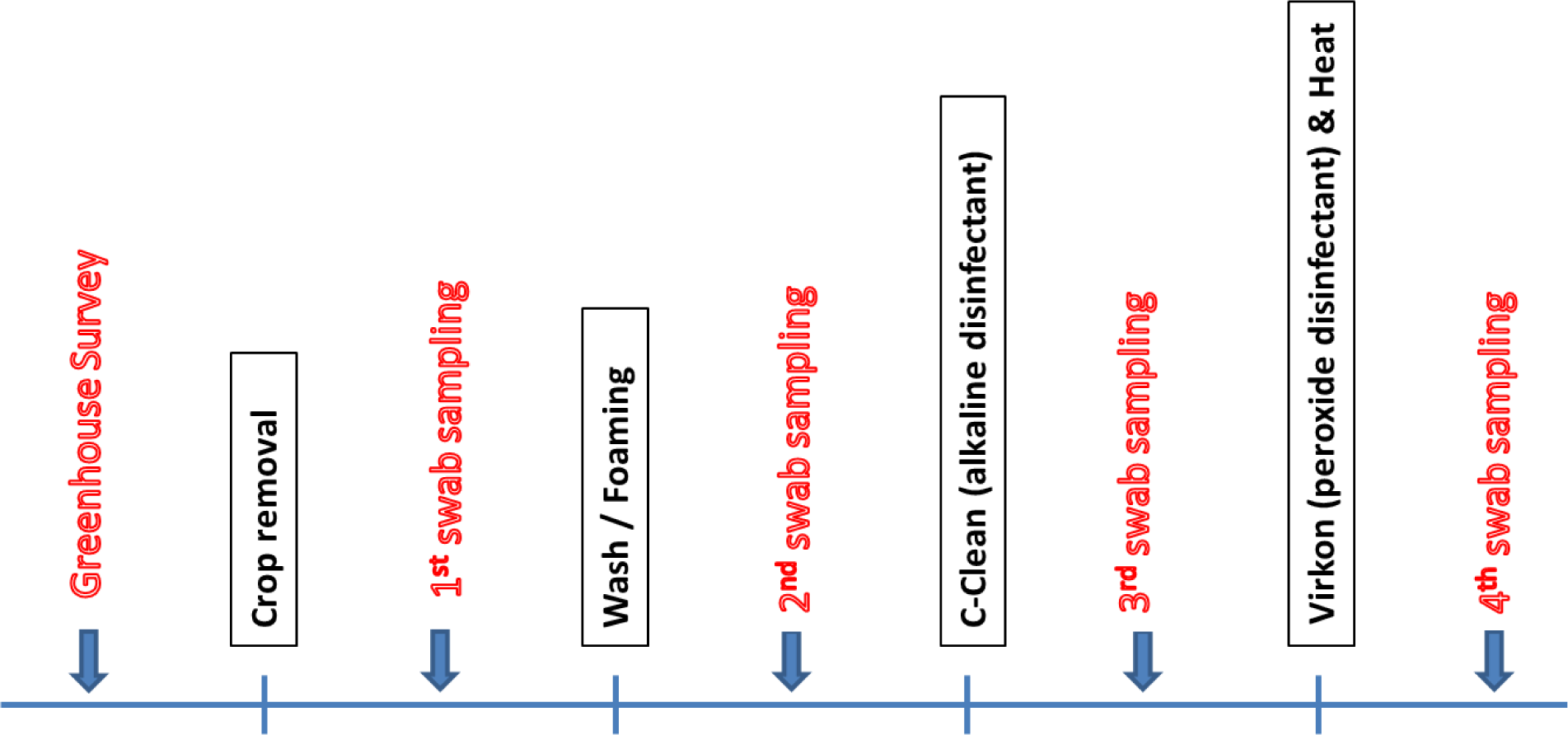
Flow chart of environmental sampling steps used in a commercial greenhouse for evaluating the effectiveness of eliminating cucumber green mottle mosaic virus using different sanitization procedures.

### Sample processing, ELISA and *in planta* virus bioassay

To facilitate extraction of liquid buffer containing the surface contaminants from the pre-moistened sampling cellulose sponges, the samples were diluted with 5 mL of sterile distilled water followed by homogenization in a Stomacher Lab Blender 400 (AJ Seward, Edmunds, UK) for 2 min. A 4.5 mL sample of the extract was used in a bioassay to confirm the presence and infectivity of the virus following the procedure described below. The remaining portion of the sample was analyzed for CGMMV particles via quantification using the CGMMV-ELISA kit (Agdia Inc., Elkhart, IN, USA) according to the manufacturer’s instructions. ELISA plates were read at OD_405 nm_ with a ‘Synergy HT’ microplate reader using ‘GEN5™’ software version 2.04. (Bio Tek^®^ Instruments Inc., Winooski, VT, USA). Negative (liquid buffer from the Solar-Cult^®^ Pre-moistened Sampling Cellulose Sponge Kit) and positive (crude extracts from cucumber leaves infected with CGMMV) controls were run in triplicate on each plate. A sample was considered CGMMV-infected if its OD_405nm_ absorbance value was at least two times greater than the negative control.

To confirm the presence and infectivity of the virus, an *in planta* bioassay was conducted by mechanically inoculating three susceptible mini cucumber seedlings of the variety ‘Picowell’ at the four true-leaf stage with the extract from each cellulose sponge. Mechanical inoculation was performed by rubbing the surface of the leaf with scrub sponge soaked in inoculum in such a way as to break the surface cells without causing too much leaf damage. Negative controls consisted of non-inoculated plants and the positive control was inoculated with CGMMV. CGMMV-infected plants first displayed symptoms at 18 d post-inoculation (dpi). Symptomatic and asymptomatic cucumber plants were tested 40 dpi to confirm the presence or absence of CGMMV using the CGMMV ImmunoStrip^®^ test kit (Agdia, Inc., Elkhart, IN) according to the manufacturer’s instructions.

### Evaluation of commercial cucumber varieties for their resistance to CGMMV and yield potential

Six Mini and nine Long English (LE) cucumber varieties used by commercial cucumber growers and with different levels of resistance to CGMMV (based on the seed companies data) and other pathogens were screened for their resistance potential to CGMMV and effects of infection on cucumber plant productivity (Table 2). Only certified healthy cucumber seeds and seedlings were used in this experiment. Seeds were germinated in rockwool cubes (Cultilene Pc, Saint-Gobain, The Netherlands) placed on flood tables in a propagation house. Rockwool cubes were soaked in water or nutrient solution for 5 min daily. Seedlings were grown at day/night temperatures of 25/20 ± 2 °C and 16 h photoperiod for 15 d prior to transplanting. Cucumber plants were transplanted at the four-leaf stage and cultivated under optimal commercial fertilization and environment conditions for 12 wk during the 2015 summer season. The experiment was conducted in one multi-span 750 m^2^ glass greenhouse compartment in the Greenhouse Research and Production Complex at the Crop Diversification Centre South, Brooks, Alberta. Coconut Coir Growbags (Millenniumsoils Coir™, St. Catharines, ON) were used to support hydroponic production of high-wire cucumber crops on raised-troughs. All the treatments were arranged in a randomized complete block design with three replicates of each of the treatments. Each treatment contained 24 cucumber plants planted in four coconut coir growbags and organized in one row. This resulted in a plant density of 3.2 plants/sq m. The nutrient solution of same standard composition was supplied to cucumber plants via a fertigation system using irrigation drippers. Plants received 2 L/h of nutrient solution containing ppm of 200 N, 55 P, 350 K, 220 Ca, 135 S, 75 Mg, 4 Fe, 0.8 Mn, 0.6 B, 0.2 Cu, 0.4 Zn and 0.17 Mo. The corresponding electrical conductivity (EC) of the nutrient solution was 2 ± 0.2 dS/m. Irrigation management and climate set points, such as temperature (day/night 23/18 ± 2 °C), light (20 h photoperiod), CO_2_ (900 µmol/mol) and relative humidity (0.5 KPa Vapor Pressure Deficit) were controlled through an automated Argus computer control system (Argus Control Systems Limited, Surrey, BC).

**Table 2.** Assessment of residual Cucumber green mottle mosaic virus (CGMMV) using enzyme-linked immunosorbent assay (ELISA) on various surfaces post treatments with various cleaning and sanitization procedures in a commercial greenhouse.

|  | Post-crop<br>removal | Post-MS<br>Topfoam<br>cleansing | Post-C-Clean<br>disinfection | Post-Virkon &<br>Heat<br>disinfection |
| --- | --- | --- | --- | --- |
| Tray top | **0.303 ± 0.019<sup>a</sup>** | 0.078 ± 0.005<sup>b</sup> | 0.078 ± 0.004<sup>b</sup> | 0.073 ± 0.001<sup>b</sup> |
| Tray gutter | **0.308 ± 0.035<sup>a</sup>** | **0.127 ± 0.026<sup>b</sup>** | 0.120 ± 0.024<sup>b</sup> | 0.094 ± 0.015<sup>b</sup> |
| Cement alleyway | **0.292 ± 0.039<sup>a</sup>** | **0.149 ± 0.029<sup>b</sup>** | **0.131 ± 0.030<sup>b</sup>** | 0.078 ± 0.005<sup>b</sup> |
| Floor mats | **0.210 ± 0.027<sup>a</sup>** | **0.152 ± 0.044<sup>ab</sup>** | 0.095 ± 0.010<sup>b</sup> | 0.074 ± 0.001<sup>b</sup> |
| Rails | **0.259 ± 0.033<sup>a</sup>** | 0.095 ± 0.010<sup>b</sup> | 0.095 ± 0.019<sup>b</sup> | 0.097 ± 0.023<sup>b</sup> |
| Rail brackets | **0.296 ± 0.045<sup>a</sup>** | 0.119 ± 0.020<sup>b</sup> | 0.101 ± 0.012<sup>b</sup> | 0.117 ± 0.028<sup>b</sup> |
| Support posts | **0.186 ± 0.020<sup>a</sup>** | 0.087 ± 0.005<sup>b</sup> | 0.085 ± 0.011<sup>b</sup> | 0.081 ± 0.009<sup>b</sup> |
| Water hoses below trays | **0.231 ± 0.020<sup>a</sup>** | 0.101 ± 0.022<sup>b</sup> | 0.094 ± 0.012<sup>b</sup> | 0.079 ± 0.007<sup>b</sup> |
| Rear tray support | **0.189 ± 0.070<sup>a</sup>** | 0.088 ± 0.011<sup>a</sup> | 0.082 ± 0.006<sup>a</sup> | 0.089 ± 0.008<sup>a</sup> |
| Canopy heating pipes | **0.139 ± 0.018<sup>a</sup>** | 0.093 ± 0.009<sup>b</sup> | 0.114 ± 0.009<sup>ab</sup> | 0.078 ± 0.005<sup>b</sup> |
| Rear end irrigation hoses | **0.137 ± 0.034<sup>a</sup>** | 0.101 ± 0.022<sup>a</sup> | 0.096 ± 0.012<sup>a</sup> | 0.093 ± 0.004<sup>a</sup> |
| Interior walls | 0.086 ± 0.005<sup>a</sup> | 0.087 ± 0.008<sup>a</sup> | 0.079 ± 0.005<sup>a</sup> | 0.080 ± 0.006<sup>a</sup> |
| Perimeter heating pipes | 0.107 ± 0.003<sup>a</sup> | 0.083 ± 0.003<sup>b</sup> | 0.087 ± 0.009<sup>b</sup> | 0.077 ± 0.003<sup>b</sup> |
| Walls-shade curtain | 0.074 ± 0.001<sup>a</sup> | 0.081 ± 0.006<sup>a</sup> | 0.080 ± 0.007<sup>a</sup> | 0.072 ± 0.000<sup>a</sup> |
| Cropping wires | 0.109 ± 0.015<sup>a</sup> | 0.080 ± 0.004<sup>a</sup> | 0.092 ± 0.017<sup>a</sup> | 0.095 ± 0.011<sup>a</sup> |
| Negative control | 0.075 ± 0.001<sup>a</sup> | 0.071 ± 0.001<sup>ab</sup> | 0.069 ± 0.001<sup>b</sup> | 0.073 ± 0.001<sup>a</sup> |
| Positive control | **0.346 ± 0.002<sup>a</sup>** | **0.341 ± 0.001<sup>b</sup>** | **0.346 ± 0.001<sup>a</sup>** | **0.348 ± 0.001<sup>a</sup>** |

\* OD<sub>405nm</sub> absorbance value ± standard error indicative of levels of residual virus contamination and effect of different cleaning/disinfecting steps on 15 different hard surfaces in a commercial cucumber greenhouse. Samples with OD<sub>405nm</sub> absorbance values in bold were considered to be CGMMV infected (positive) and virus infectivity was confirmed in a greenhouse bioassay. Least square means are not significantly different according to ANOVA LSMeans Student's *t* tests when followed by the same letter on the same row ( $\alpha = 0.05$ , $n = 3$ ).

The crop was monitored for insects, mites and diseases during the season using standard integrated pest management (IPM) scouting techniques. Two Floramite^®^ SC (bifenazate 22.6% SU) (Chemtura Canada Co./Cie, Elmira, ON), two Dibrom^®^ (naled 900g/L EC) (Loveland Products Canada Inc., Dorchester, ON) and one Kopa Insecticidal Soap (potassium salts of fatty acids 47.0% SN) (Neudorff North America, Saanichton, BC) spray applications were required during the crop trial for the control of spider mites.

### Inoculum collection and maintenance

The CGMMV strain used as inoculum in this study was isolated from a diseased cucumber plant collected in a commercial greenhouse in Alberta, Canada and maintained through rub-inoculation on the susceptible mini cucumber variety ‘Picowell’ in a growth chamber. The complete genome sequencing of this CGMMV isolate (GenBank accession no. KP772568) revealed that it was closely related by 98% to 99% nucleotide sequence identities to isolates of Asian origin (Li *et al.*, 2015).

### Monitoring the presence and spreading pattern of CGMMV in a greenhouse

The primary source of infection was introduced to the greenhouse through the inoculation of the first plant of each row with the CGMMV strain 7 d after transplanting (4 true-leaf stage) through rub-inoculation, as described above (Fig. 1). The secondary spread of the virus to the neighboring plants occurred mechanically through plant handling, leaf contact, wounds made from cutting tools, or farming equipment. Cucumber plants were twisted, clipped, de-leafed and lowered twice/wk and fruits were harvested thrice/wk. Before moving from one plot to the next, new gloves were worn and the cutting tools and farming equipment were disinfected using 2% Virkon (potassium peroxymonosulphate 21.4% SP).

All cucumber plants were visually examined weekly for the appearance of CGMMV symptoms, starting one-week post-inoculation. Symptomatic and asymptomatic cucumber plants were tested to confirm the presence or absence of CGMMV using the ImmunoStrip^®^ test kit.

### Data collection and statistical analyses

Disease incidence was recorded weekly. Area under disease progress curve (AUDPC) was calculated for each plot according to Simko and Piepho (2012) as follows:

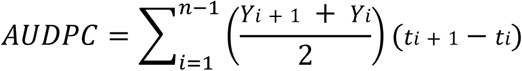

where *Y*_*i*_ = the proportion of diseased cucumber plants at *i*th observation, *t*_*i*_ = time of the *i*th observation in days from the first observation and *n* = total number of disease observations. AUDPC was used to assess quantitative disease resistance in cucumber varieties (Jeger and Viljanen-Rollinson, 2001).

Cucumber fruits were harvested thrice/wk and the number of fruit and fruit weight/plot were recorded. The data was subjected to ANOVA and, in the presence of treatment effects, the statistical significance of differences between treatments means was assessed using the Least Significant Difference (LSD) test.

### Evaluation of grafted cucumber plants for resistance to CGMMV and yield potential

Grafted plants consisted of scion and rootstock of the CGMMV susceptible Mini cucumber variety ‘Picowell’ and the CGMMV highly resistant (seed company data) LE cucumber variety ‘Bonbon’, respectively. Certified healthy cucumber seeds were germinated in Cultilene Pc rockwool cubes placed on flood tables in a propagation house. Rockwool cubes were soaked in water or nutrient solution for 5 min daily. Seedlings were grown at 25/20 ± 2 °C day/night and 16 h photoperiod for 5 d before grafting or transplanting for non-grafted plants. Grafting was conducted at the one cotyledon stage when the cotyledons of the scions and rootstocks were completely unfolded (Hassell *et al.*, 2008). The grafted seedlings were then placed in a greenhouse mist chamber, where a fine mist was delivering every 10 secs/min. Seedlings were kept in the mist chamber for 3 d, then transferred to flood tables in propagation house and grown for an additional week prior to transplanting into a poly greenhouse compartment. Grafted and non-grafted cucumber plants were transplanted at the four-leaf stage and cultivated under optimal commercial fertilization and environment conditions, as described above, for 12 wk during the fall season. Coconut coir growbags were used to support hydroponic production of the cucumber crop maintained in an umbrella system. All the treatments were arranged in a randomized complete block design with six replicates for each of the treatments. Each treatment contained one cucumber plants planted in a coconut coir growbag. Plants were either inoculated with CGMMV or sterile water (negative control) 7 d after transplanting through rub-inoculation. Weekly fruit yield and internode number were recorded. The data were subjected to ANOVA and, in the presence of treatment effects, the significance of differences between treatment means was assessed using the Least significant difference (LSD) test.

## Results

### Evaluation of the effectiveness of the different sanitization steps performed in a commercial greenhouse

There was a high variability of CGMMV frequency in all surfaces tested (Table 2). The majority of the surfaces sampled tested positive for CGMMV, with the exception of interior walls, perimeter heating pipes, walls-shade curtains and crop support wires. Surfaces directly in contact with the plants were the most heavily infested areas, with tray tops (0.303 ± 0.019) and tray gutters (0.308 ± 0.035) having the highest levels of CGMMV. Additional areas with high levels of CGMMV included rails (0.259 ± 0.033), rail brackets (0.296 ± 0.045), water hoses below trays (0.231 ± 0.020) and cement alleyway (0.292 ± 0.039). Cleansing using pressure washing and MS Topfoam reduced detectable levels of CGMMV in all treatments. However, tray tops, rails and rails brackets surfaces were easier to clean than cement alleyway, tray gutter and floor mats (Table 2). Cleansing using MS TopFoam LC Fresh cleaner (sodium hydroxide 7%, 2-(2-butoxyethoxy) ethanol 7%, tetrasodium EDTA 6%, sodium laurel sulphate 5%),was sufficient to eliminate CGMMV on all infested surfaces, except tray gutters, cement alleyways and floor mats. Two sanitization steps using MS Topfoam and C-Clean (chrorinated alkaline cleaner) were necessary to eliminate CGMMV from tray gutters and floor mats. Cement alleyway was the most difficult surface to disinfect and all cleaning/disinfecting steps were needed to eliminate CGMMV infectivity.

### Spread pattern of CGMMV infection in a greenhouse varietal screening trial

Overall, the mini cultivars of cucumbers had faster spread of CGMMV infection than Long English (LE) cultivars (Fig. 2). In mini cultivars, CGMMV symptoms first appeared at 10 dpi. There was an exponential spread of new plants showing symptoms 25 d following initial inoculation (Fig. 2). CGMMV symptom development in LE cucumbers was first observed at 10 dpi, similar to mini cultivars. However, spread of CGMMV to new plants was greatly reduced in LE cultivars as exponential spread of the virus only occurred after 52 dpi (Fig. 2). Moreover, the mini cultivars showed almost 100% infection by the end of the crop production cycle, except for the ‘Katrina’ variety which reached a maximum level of 80% infected plants. CGMMV infection of the LE cultivars was much lower ranging between 45 and 90% infected plants within the same time period. Within LE varieties, the ‘DR4879CE’ had the highest infection rate (90.28%), while ‘LC13900’, ‘Bomber’ and ‘Sepire’ all had infection levels at near 50% at the end of the crop production cycle (Fig. 2).

**Fig. 2.**
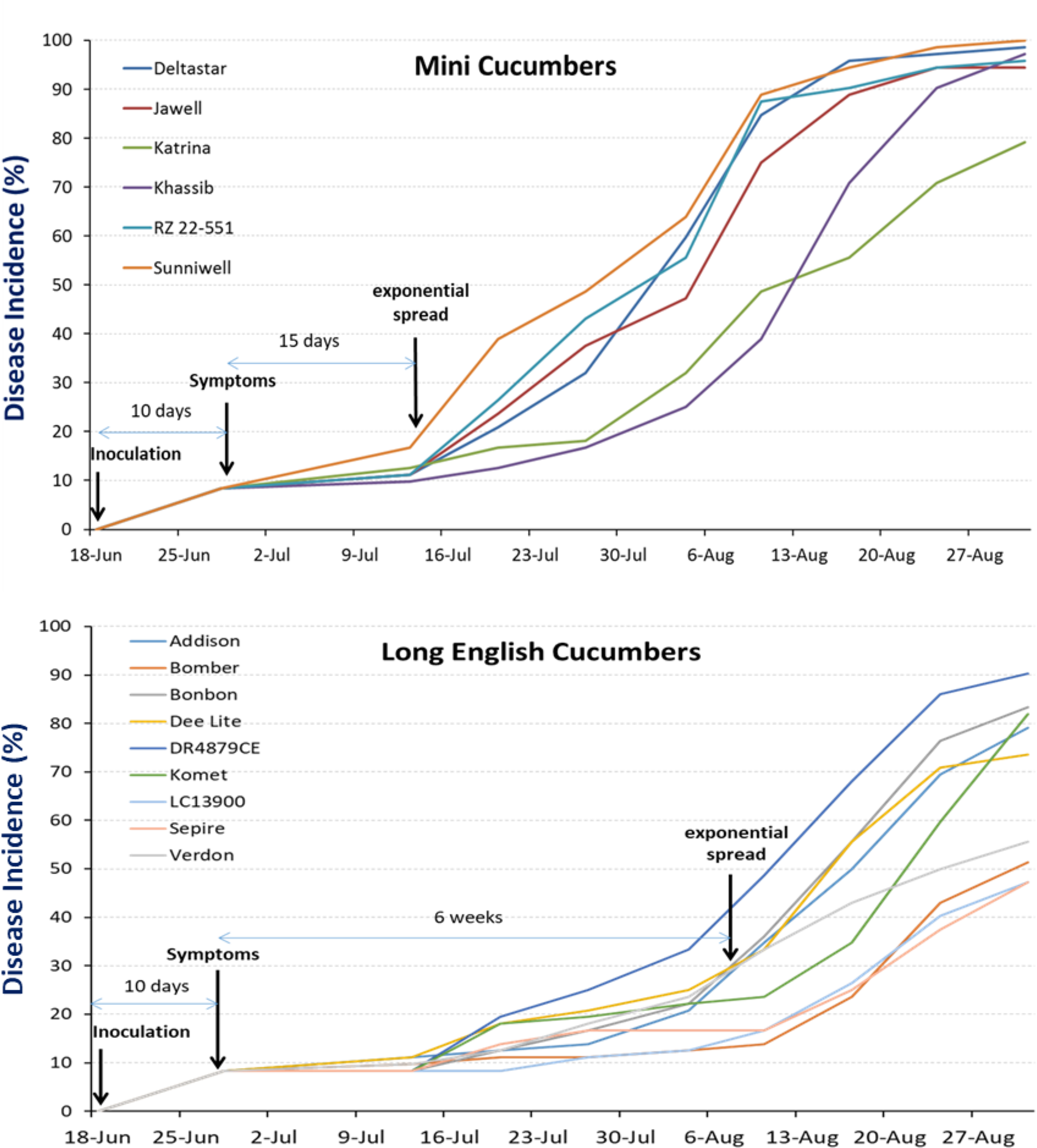
Disease progression curves of Cucumber green mottle mosaic virus as measured on various varieties of Mini and Long English types of cucumber in a crop production season.

### Levels of resistance of greenhouse cucumber varieties to CGMMV and effects of infection on productivity

Among six Mini varieties, ‘Sunniwell’ was the most tolerant to CGMMV. This variety showed high CGMMV infection level (AUDPC = 3621.53 ± 267.26), however, it had the highest fruit yield (182 ± 14.90 fruit/m^2^). ‘Katrina’ was the most resistant (AUDPC = 2163.89 ± 128.65), but was intermediate in yield compared to ‘Sunniwell’ (166.01 ± 17.96 fruit/m^2^) (Fig. 3).

**Fig. 3.**
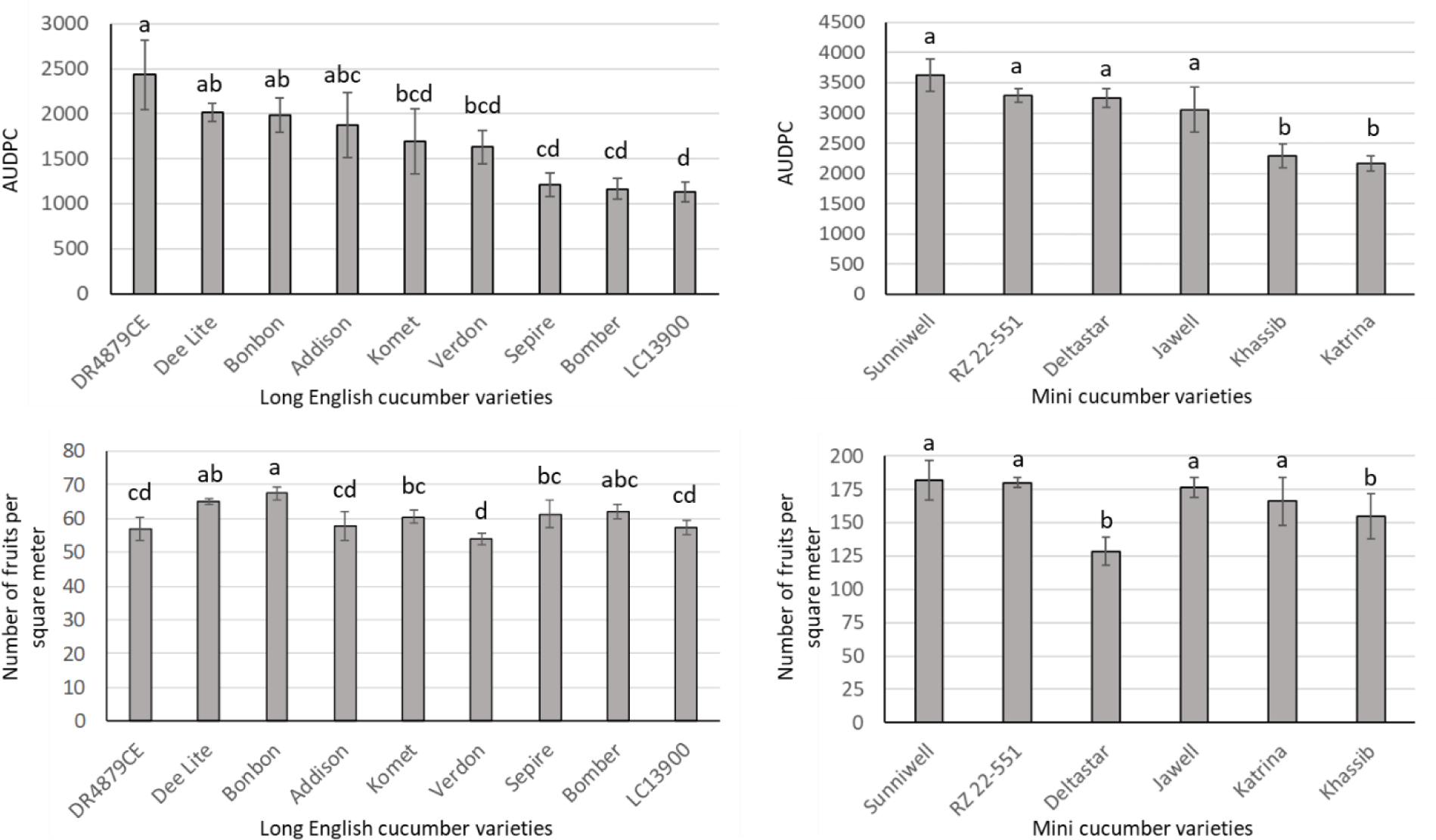
Quantitative disease resistance measured as area under disease progression curve (AUDPC) and yield in Mini and Long English greenhouse cucumber varieties infected by Cucumber green mottle mosaic virus. Bars represent least square means that are significantly different when associated with different letters, according to ANOVA LSMeans Student’s *t* tests (*α* = 0.05). Error bars indicates standard error from three repeats (*n* = 3).

Among nine varieties of LE screened for resistance to CGMMV, ‘Bonbon’ was tolerant. This variety showed a high CGMMV infection level (AUDPC = 1980.56 ± 117.98) without compromising yield, which was highest (67.42 ± 1.80 fruit/m^2^) (Fig. 3). The most susceptible LE variety was ‘DR4879CE’ (AUDPC = 2430.56 ± 389.50), while ‘Verdon’, the most widely grown cultivar in Alberta, was intermediate (AUDPC = 1629.17 ± 183.07) in its resistance (Fig. 3). In yield comparisons, ‘DR4879CE’ and ‘Verdon’ were poor performers (Fig. 3).

### Evaluation of the potential commercial viability of grafted cucumber plants for resistance to CGMMV and yield potential

In grafting experiments, the CGMMV susceptible Mini cucumber variety ‘Picowell’ was used as the scion and the LE cucumber variety ‘Bonbon’, described by the supplier as highly resistant to CGMMV, was chosen as the rootstock. The grafted plant was compared with the non-grafted counterparts for resistance to CGMMV and yield assessment. The grafted ‘Picowell’ on ‘Bonbon’ yielded 16% more crops than the non-grafted ‘Picowell’ did (Fig. 4). However, in the CGMMV infected grafts between ‘Picowell’ and ‘Bonbon’, the yield was compromised. The yield of the grafted plants was reduced to approximately 72% as compared to the non-infected grafted cucumbers. Similar yield reduction was obtained for the non-grafted cucumbers (Fig. 4). The non-grafted ‘Picowell’ produced the highest internode number and the tallest plants in both inoculated and control treatments (Fig. 4).

**Fig. 4.**
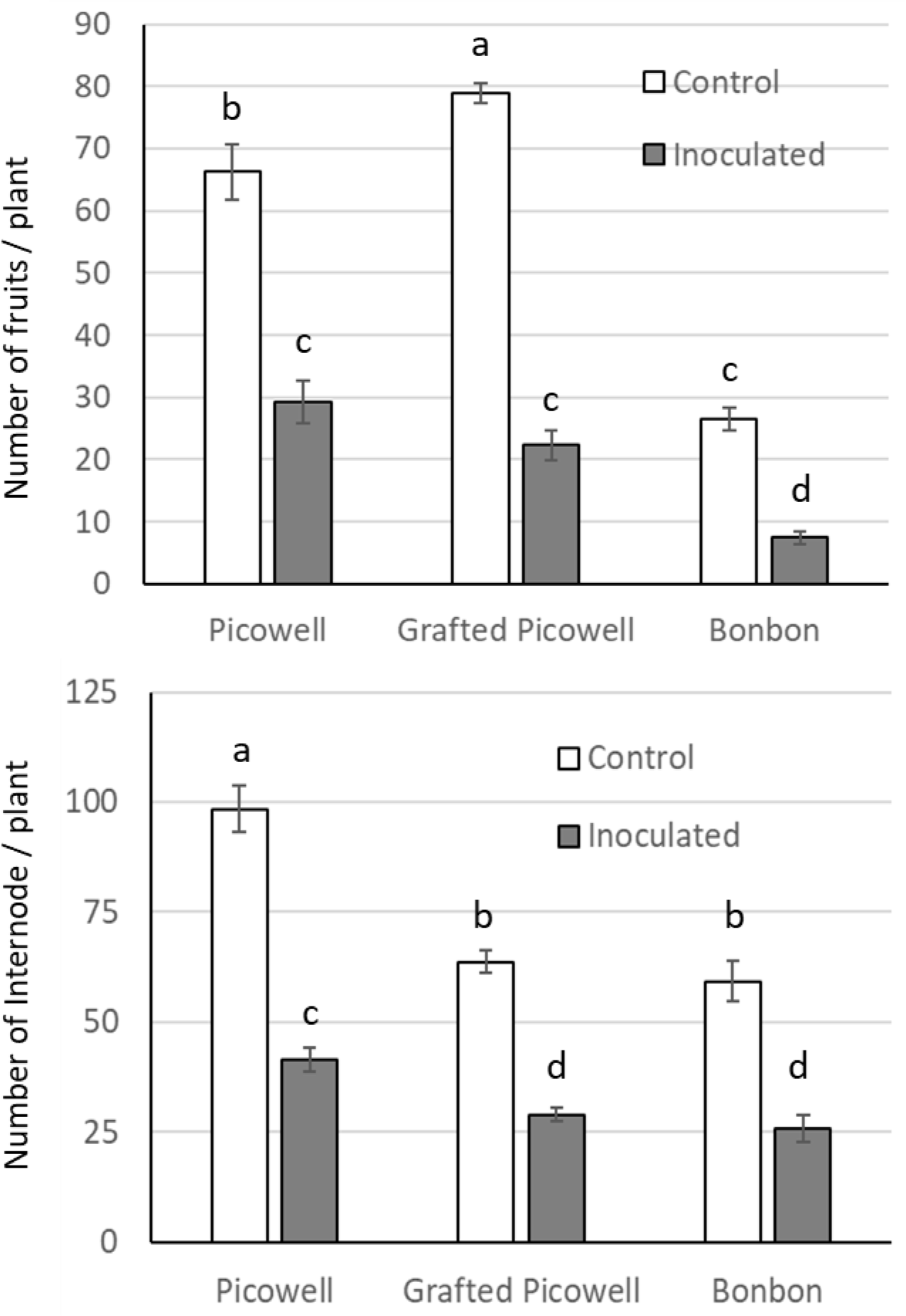
Effect of Cucumber green mottle mosaic virus infection on yield and internode number of grafted and non-grafted Mini and Long English greenhouse cucumber varieties. Bars represent least square means that are significantly different when associated with different letters, according to ANOVA LSMeans Student’s *t* tests (α = 0.05). Error bars indicates standard error from six repeats (*n* = 6).

## Discussion

CGMMV posts a serious threat to greenhouse cucumber, as well as other cucurbit crop productions worldwide. This virus is capable of rapidly spreading throughout a greenhouse or other protected cultivations, destroying crops and resulting in significant economic losses to growers (Reingold *et al.*, 2016). CGMMV can also persist in the environment, infecting new crops and perpetuating the disease cycle. To our knowledge this is the first systematic analysis of the distribution of CGMMV within a commercial greenhouse, and the first study to evaluate different steps of a decontamination approach for CGMMV in total greenhouse cleaning.

Our results demonstrated the importance of using pressure washing and cleansing with an alkaline foam cleanser as a first step in the sanitization procedure to significantly reduce or even eliminate CGMMV contamination of greenhouse surfaces. This step eliminated CGMMV on some of the most heavily infested areas and reduced detectable amounts of virus contamination on cement alleyways, tray gutters and floor mats. The critical importance of this first step of sanitization lies in the fact that it was vitally important to remove the organic matter, a primary source of disease-causing plant pathogens. This step should be carried out in advance of disinfection since some disinfectants are inactivated by direct contact with organic matter. The wide variation in antiviral activity occurring with the same sanitizer on different surfaces could be partly explained by differences in surfaces porosities (Stedman *et al.*, 1955a). Our experimental data revealed that the second and third steps of sanitization, with alkaline and peroxide disinfectants, were essential to completely eliminate CGMMV on porous and uneven surfaces, such as cement alleyways, tray gutters and floor mats. These results corroborate previous findings suggesting that, in general, all disinfectants require higher concentrations and long exposure times to reduce significantly the microbial populations on porous surfaces (Stedman *et al.*, 1955b). Furthermore, it was found that two successive applications of disinfectant were more effective than a single, prolonged application in most instances due to a residual disinfectant activity remaining from the previous treatment (Stedman *et al.*, 1955b).

The host resistance response to viral infection is often the most important aspect of control. Out of 15 cucumber varieties evaluated, two Mini and three LE had reduced or delayed CGMMV infection spread in the greenhouse. These varieties were similar to a group of cucumber cultivars reported by Čech and Branišová (1976) that consisted of plants with paradoxically high symptomatic sensitivity and high resistance to virus increase. Cucumber plants from this group were inoculated with a stock virus suspension and many of them failed to become infected. The reduced spread of CGMMV may be associated with low viral titer accumulation due to low virus reproduction rate in these varieties. A recent study identified two partially resistant *C. sativus* accessions out of 58 evaluated using quantitative RT-PCR (Crespo *et al.*, 2018). These accessions showed only mild CGMMV disease symptoms and low viral titer accumulation. Cucumber varieties with low virus replication capacity within the plant may well be excellent candidates for breeding programs developing CGMMV-resistant varieties.

The use of high-yielding mini and LE cucumber varieties that can delay CGMMV spread, as demonstrated in this trial, is recommended for cultivation in the greenhouses assuming that this would reduce the incidence and spread of CGMMV infection and could reduce economic losses to cucumber growers. However, growers should use efficient sanitization programs between crops, since repeated production with partially resistant cultivars may increase the level of CGMMV particles on various hard surfaces within the greenhouse rendering even partial resistance ineffective. Hence the urgent need for resistant cultivars with restricted virus movement and replication.

The spread of CGMMV had no specific trends (S.1), revealing that cultural practices (pruning, de-leafing, lowering, fruit picking) were not the only factors responsible for CGMMV infection spread. Other factors, such as leachates, chewing insects, bumble bees and contaminated seeds, were possibly playing a role in the spread of the infection. In the present experiment, bumble bees were not used for cucumber plant pollination. However, it is possible that bumble bees moved from an adjacent tomato crop and inadvertently contributed to the CGMMV infection spread between cucumber plants. The contribution of honey bees to the spread of CGMMV infection was previously demonstrated by Darzi *et al.* (2018). The control of pollinator insects would reduce the incidence and spread of CGMMV infection between cucumber plants.

Our trial revealed that grafting did not increase yields of the CGMMV-infected grafted plants between ‘Picowell’ and ‘Bonbon’, as compared to the CGMMV infected non-grafted ‘Picowell’. However, yield enhancement was successfully achieved using the same scion/rootstock combination in CGMMV disease-free environment. In previous studies, rootstocks have been observed to increase or decrease the incidence of non-soilborne virus infection in the scion (Davis *et al.*, 2008). In our greenhouse varietal screening trial, ‘Bonbon’ showed high CGMMV infection levels. It is possible that the level of resistance of ‘Bonbon’ to the Canadian CGMMV isolate was not sufficient to confer CGMMV resistance to the scion. It would be interesting to determine if the use of rootstocks with higher level of resistance to CGMMV, such as ‘Sepire’, ‘Bomber’ and ‘LC13900’, would confer virus resistance to the scion.

Results of the present study are in agreement with the previous studies reporting the positive effect of grafting on fruit yield (Davis *et al.*, 2008; Usanmaz and Abak, 2019). Seong *et al.* (2003) reported 27% increases in the marketable yield of cucumbers from grafted plants when compared to the non-grafted scion cultivars. The higher fruit yield and reduced shoot growth of grafted cucumber plants reported in the present study are possibly due to the redirection of assimilates away from vegetative growth toward reproductive organs.

## Conclusions

Our study highlighted the importance of pressure washing and cleansing as a first step of the sanitization procedure in eliminating CGMMV on some of the most heavily infested areas. However, second and third steps using different disinfectants were essential to eliminate CGMMV on porous and uneven surfaces. These results are key in developing efficient methods for decontamination of commercial greenhouses infested with CGMMV.

The varietal screening trial for resistance to CGMMV revealed the relative suitability of commercial cucumber varieties for use in greenhouses at risk from CGMMV infection and where minimizing production losses is a key consideration. While some varieties can restrain CGMMV spread, most varieties tested showed susceptibility to CGMMV demonstrating the need for new productive cucumber varieties with CGMMV resistance.

Grafting cucumber plants have been reported in this study to increase fruit yield in CGMMV disease-free environment. However, grafting did not increase yields of the CGMMV-infected grafts. Further testing of different scion/rootstock combinations is needed to determine if the use of rootstocks with higher level of resistance to CGMMV will confer virus resistance to the scion without compromising the fruit yield.

## Acknowledgements

The authors gratefully acknowledge the financial support of the Alberta Crop Industry Development Fund (grant numbers: 2014C005R and 2016C007R) and the Business Development and Farm Safety Internal Initiatives through Growing Forward 2, a Federal-Provincial-Territorial Initiative (grant number: 609126). Technical assistance provided by members of the Greenhouse Section at the Crop Diversification Centre South, Brooks, AB is sincerely appreciated. Thanks are also due to Dr. Jonathan Griffiths and Dr. Antonet Svircev, Agriculture and Agri-Food Canada, Vineland Station, ON for their helpful comments on the manuscript.

## References

Ainsworth, G.C., 1935. Mosaic diseases of the cucumber. Annals of Applied Biology 22, 55–67.

Al-Shahwan, I.M., Abdalla, O.A., 1992. A Strain of Cucumber green mottle mosaic virus (CGMMV) from bottlegourd in Saudi Arabia. Journal of Phytopathology 134, 152–156.

Borodynko-Filas, N., Minicka, J., Hasiów-Jaroszewska, B., 2017. The Occurrence of Cucumber green mottle mosaic virus infecting greenhouse cucumber in Poland. Plant Disease 101, 1336–1336.

Boubourakas, I.N., Hatziloukas, E., Antignus, Y., Katis, N.I., 2004. Etiology of leaf chlorosis and deterioration of the fruit interior of watermelon plants. Journal of Phytopathology 152, 580–588.

Budzanivska, I.G., Rudneva, T.O., Shevchenko, T.P., Boubriak, I., Polischuk, V.P., 2007. Investigation of Ukrainian isolates of cucumber green mottle mosaic virus. Archives of Phytopathology and Plant Protection 40, 376–380.

Cech, M., Branišová, H., 1976. Nonrelatedness between symptoms and cucumber virus 4 content in different cucumber cultivars. Biologia plantarum 18, 58–62.

Celix, A., Luis-Arteaga, M., Rodriguez-Cerezo, E., 1996. First report of Cucumber green mottle mosaic *Tobamovirus* infecting greenhouse-grown cucumber in Spain. Plant Disease 80, 1303.

Crespo, O., Janssen, D., Robles, C., Ruiz, L., 2018. Resistance to Cucumber green mottle mosaic virus in *Cucumis sativus*. Euphytica 214, 201.

Darzi, E., Smith, E., Shargil, D., Lachman, O., Ganot, L., Dombrovsky, A., 2018. The honeybee *Apis mellifera* contributes to Cucumber green mottle mosaic virus spread via pollination. Plant Pathology 59, 244–251.

Davis, A.R., Perkins-Veazie, P., Sakata, Y., López-Galarza, S., Maroto, J.V., Lee, S.-G., Huh, Y.-C., Sun, Z., Miguel, A., King, S.R., Cohen, R., Lee, J.-M., 2008. Cucurbit grafting. Critical Reviews in Plant Sciences 27, 50–74.

Dombrovsky, A., Tran-Nguyen, L.T.T., Jones, R.A.C., 2017. Cucumber green mottle mosaic virus: rapidly increasing global distribution, etiology, epidemiology, and management. Annual Review of Phytopathology 55, 231–256.

Ellouze, W., Dalpé, S., Howard, R.J., Ling, K.-S., Zhang, W., 2015. Managing Cucumber green mottle mosaic virus in Alberta greenhouses, Agri-Facts: Practical Information for Alberta’s Agriculture Industry. Alberta Agriculture and Forestry, http://www1.agric.gov.ab.ca/$department/deptdocs.nsf/all/agdex15624.

Fletcher, J.T., George, A.J., Green, D.E., 1969. Cucumber green mottle mosaic virus, its effect on yield and its control in the Lea Valley, England. Plant Pathology 18, 16–22.

Gu, J.T., Fan, S.X., Zhang, X.C., 2008. Effects of rootstocks on the development, disease resistance and quality of *Cucumis sativus* L. Acta Horticulturae 771, 161–166.

Guan, W., Zhao, X., Hassell, R., Thies, J., 2012. Defense mechanisms involved in disease resistance of grafted vegetables. HortScience 47, 164–170.

Hasama, W., Morita, S., Kato, T., 1993. Reduction of resistance to corynespora target leaf spot in cucumber grafted on a bloomless rootstock. Japanese Journal of Phytopathology 59, 243–248.

Hassell, R.L., Memmott, F., Liere, D.G., 2008. Grafting methods for watermelon production. HortScience 43, 1677–1679.

Jeger, M.J., Viljanen-Rollinson, S.L.H., 2001. The use of the area under the disease-progress curve (AUDPC) to assess quantitative disease resistance in crop cultivars. Theoretical and Applied Genetics 102, 32–40.

Kim, O.-K., Mizutani, T., Natsuaki, K.T., Lee, K.-W., Soe, K., 2010. First report and the genetic variability of Cucumber green mottle mosaic virus occurring on bottle gourd in Myanmar. Journal of Phytopathology 158, 572–575.

Lecoq, H., Desbiez, C., 2012. Viruses of cucurbit crops in the mediterranean region: an ever-changing picture, In: Loebenstein, G., Lecoq, H. (Eds.), Advances in Virus Research. Academic Press, pp. 67–126.

Li, J.-X., Liu, S.-S., Gu, Q.-S., 2016. Transmission efficiency of Cucumber green mottle mosaic virus via seeds, soil, pruning and irrigation water. Journal of Phytopathology 164, 300–309.

Li, R., Zheng, Y., Fei, Z., Ling, K.-S., 2015. First complete genome sequence of an emerging Cucumber green mottle mosaic virus isolate in North America. Genome Announcements 3.

Ling, K.-S., Li, R., Zhang, W., 2014. First report of Cucumber green mottle mosaic virus infecting greenhouse cucumber in Canada. Plant Disease 98, 701–701.

Liu, H.W., Luo, L.X., Li, J.Q., Liu, P.F., Chen, X.Y., Hao, J.J., 2014. Pollen and seed transmission of Cucumber green mottle mosaic virus in cucumber. Plant Pathology 63, 72–77.

Louws, F.J., Rivard, C.L., Kubota, C., 2010. Grafting fruiting vegetables to manage soilborne pathogens, foliar pathogens, arthropods and weeds. Scientia Horticulturae 127, 127–146.

Mandal, S., Mandal, B., Mohd, Q., Haq, R., Varma, A., 2008. Properties, diagnosis and management of Cucumber green mottle mosaic virus. Plant Virus 2, 25–34.

Melnyk, C.W., 2016. Plant grafting: insights into tissue regeneration. Regeneration (Oxf) 4, 3–14.

Moradi, Z., Jafarpour, B., 2011. First report of coat protein sequence of Cucumber green mottle mosaic virus in cucumber isolated from Khorasan in Iran. International Journal of Virology 7, 1–12.

Pérez-Alfocea, F., 2015. Why should we investigate vegetable grafting?, Acta Horticulturae 1086: I International Symposium on Vegetable Grafting ed. International Society for Horticultural Science (ISHS), Leuven, Belgium, pp. 21–29.

Rao, A.L.N., Varma, A., 1984. Transmission studies with Cucumber green mottle mosaic virus. Journal of Phytopathology 109, 325–331.

Reingold, V., Lachman, O., Belausov, E., Koren, A., Mor, N., Dombrovsky, A., 2016. Epidemiological study of Cucumber green mottle mosaic virus in greenhouses enables reduction of disease damage in cucurbit production. Annals of Applied Biology 168, 29–40.

Reingold, V., Lachman, O., Koren, A., Dombrovsky, A., 2013. First report of Cucumber green mottle mosaic virus (CGMMV) symptoms in watermelon used for the discrimination of non-marketable fruits in Israeli commercial fields, New Disease Reports, p. 11.

Rouphael, Y., Kyriacou, M.C., Colla, G., 2018. Vegetable grafting: A toolbox for securing yield stability under multiple stress conditions. Frontiers in Plant Science 8.

Sakata, Y., Sugiyama, M., Ohara, T., Morishita, M., 2006. Influence of rootstocks on the resistance of grafted cucumber (*Cucumis sativus* L.) scions to powdery mildew (*Podosphaera xanthii* U. Braun & N. Shishkoff). Journal of the Japanese Society for Horticultural Science 75, 135–140.

Seong, K.C., Moon, J.H., Lee, S.G., Kang, Y.G., Kim, K.Y., Seo, H.D., 2003. Growth, lateral shoot development, and fruit yield of white-spined cucumber (*Cucumis sativus* cv. Baekseong-3) as affected by grafting methods. Journal of the Korean Soceity for Horticultural Science 44, 478–482.

Shargil, D., Smith, E., Lachman, O., Reingold, V., Darzi, E., Tam, Y., Dombrovsky, A., 2017. New weed hosts for Cucumber green mottle mosaic virus in wild Mediterranean vegetation. European Journal of Plant Pathology 148, 473–480.

Shibuya, T., Itagaki, K., Wang, Y., Endo, R., 2015. Grafting transiently suppresses development of powdery mildew colonies, probably through a quantitative change in water relations of the host cucumber scions during graft healing. Scientia Horticulturae 192, 197–199.

Simko, I., Piepho, H.-P., 2012. The area under the disease progress stairs: calculation, advantage, and application. Analytical and Theoretical Plant Pathology 102, 381–389.

Stedman, R.L., Kravitz, E., Bell, H., 1955a. Studies on the efficiencies of disinfectants for use on inaminate objects. III. Physicochemical factors affecting survace disinfection. Applied microbiology 3, 71–74.

Stedman, R.L., Kravitz, E., Bell, H., 1955b. Studies on the efficiencies of disinfectants for use on inanimate objects. IV. Factors of importance in practical disinfecting procedures. Applied microbiology 3, 273–276.

Sui, X., Li, R., Shamimuzzaman, M., Wu, Z., Ling, K.-S., 2019. Understanding the transmissibility of Cucumber green mottle mosaic virus in watermelon seeds and seed health assays. Plant Disease 103, 1126–1131.

Tesoriero, L.A., Chambers, G., Srivastava, M., Smith, S., Conde, B., Tran-Nguyen, L.T.T., 2016. First report of Cucumber green mottle mosaic virus in Australia. Australasian Plant Disease Notes 11, 1.

Tian, T., Posis, K., Maroon-Lango, C.J., Mavrodieva, V., Haymes, S., Pitman, T.L., Falk, B.W., 2014. First report of Cucumber green mottle mosaic virus on melon in the United States. Plant Disease 98, 1163–1163.

Usanmaz, S., Abak, K., 2019. Plant growth and yield of cucumber plants grafted on different commercial and local rootstocks grown under salinity stress. Saudi Journal of Biological Sciences 26, 1134–1139.

Varveri, C., Vassilakos, N., Bem, F., 2002. Characterization and detection of Cucumber greem mottle mosaic virus in Greece. Phytoparasitica 5, 493 – 501.

Wang, J., Zhang, D., Fang, Q., 2002. Studies on antivirus disease mechanism of grafted seedless watermelon. Anhui Nongxueyuan Xuebao 29, 336–339.

Zhang, Y.J., Li, G.F., Li, M.F., 2009. Occurrence of Cucumber green mottle mosaic virus on Cucurbitaceous Plants in China. Plant Disease 93, 200–200.

