## Supplementary material for "Control of Cucumber green mottle mosaic virus in commercial greenhouse production with agricultural disinfectants and resistant cucumber varieties"

### Supplementary materials

**S.1** Disease progression and spread patterns of Cucumber green mottle mosaic virus (CGMMV) infection in Mini and Long English cucumber varietal trials, based on symptom observation and test confirmation for the CGMMV infection. Red circles represent plants inoculated with CGMMV through rub-inoculation. Hollow red circles represent plants infected with CGMMV through plants handling after touching the CGMMV inoculated plant. Numbers from 1-6 represent Mini varieties: 1. Sunniwell; 2. Deltastar; 3. RZ 22-551; 4. Khassib; 5. Jawell and 6. Katrina. Numbers from 7-15 represent Long English varieties: 7. DR4879CE; 8. Bomber; 9. LC13900; 10. Dee Lite; 11. Komet; 12. Bonbon; 13. Verdon; 14. Addison and 15. Sepire.

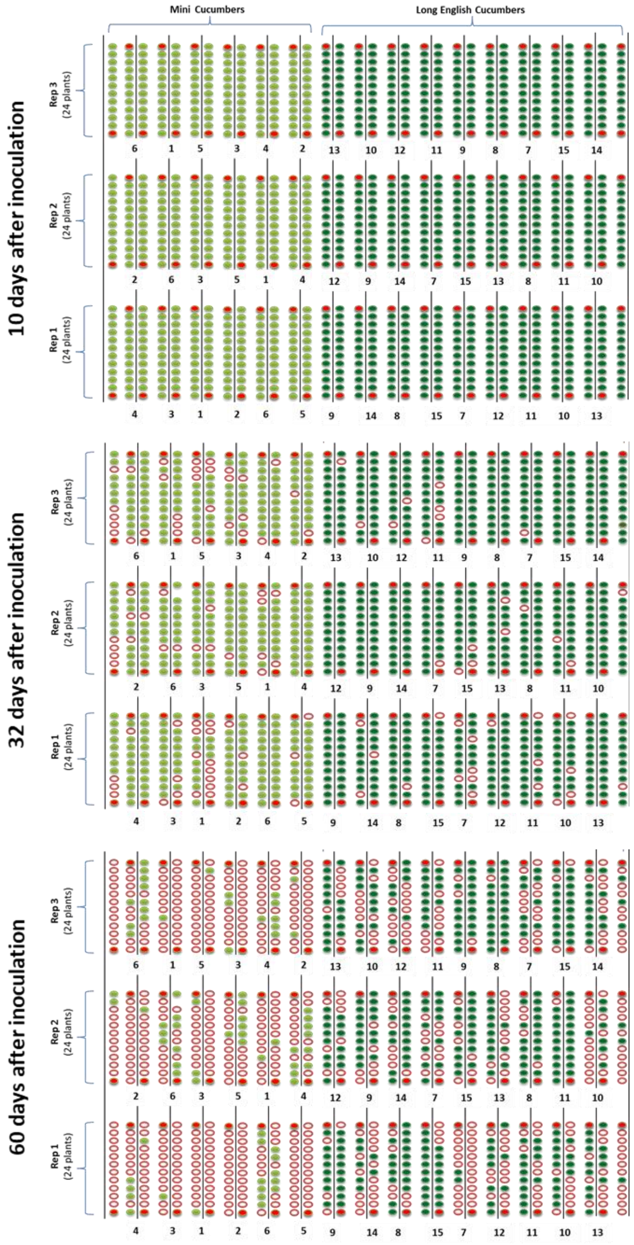
